# Environment-mediated metabolic constraints shape antimicrobial resistance evolution in *Pseudomonas aeruginosa*

**DOI:** 10.64898/2026.10.19.760112

**Authors:** Joseph M. Ficarrotta, Glynis L. Kolling, Jason A. Papin

## Abstract

*Pseudomonas aeruginosa* rapidly evolves resistance to multiple antibiotics, complicating treatment of chronic infections. While resistance mechanisms are well described, the influence of host-relevant environmental conditions on resistance evolution remains unclear. Here, we investigated how nutrient composition, pH, and oxygen availability influence resistance evolution to piperacillin, tobramycin, and ciprofloxacin. We evolved *P. aeruginosa* in different synthetic media mimicking different infection niches as well as variations in pH and oxygen availability. We also performed whole-genome sequencing to identify mutational adaptive strategies. Our findings reveal that environmental context profoundly affects both the rate and mechanisms of resistance evolution. Nutrient-rich conditions and aerobic metabolism favored canonical resistance mutations, while nutrient-limited or anaerobic environments promoted adaptations involving efflux regulation, metabolic remodeling, and biofilm-associated traits. pH and oxygen availability acted as selective pressures, modulating which pathways were accessible as well as the stability of resistance phenotypes. Additionally, the mutations that arose in the lab-evolved isolates were also actively maintained across a large population of clinical *P. aeruginosa* genomes, confirming that spatial oxygen gradients in host tissues drive parallel evolutionary pathways *in vivo*. These results underscore that resistance evolution is not uniform but highly contingent on the metabolic and redox environment. Understanding how infection-specific conditions shape evolutionary trajectories provides insight into the emergence of multidrug resistance and highlights potential avenues for therapeutic intervention.

**Importance:** Antibiotic resistance in *Pseudomonas aeruginosa* is a leading cause of treatment failure in chronic infections, yet the relationship between the infection-site environment, bacterial metabolism, and the development of resistance remains poorly understood. Here, we use synthetic media that mimic chronic wounds, urinary tract infections, and the cystic fibrosis lung to demonstrate that the development of resistance is both drug- and niche-specific, with pH and oxygen availability of further shaping evolutionary outcomes in drug-dependent manner. Whole-genome sequencing reveals that oxygen availability during evolution selects for distinct resistance mutations. These findings are recapitulated in a clinical cohort of *P. aeruginosa* genomes, confirming their translational relevance. Together, these results establish resistance evolution as a highly environment-contingent process and suggest that a deeper understanding of how infection-site conditions constrain evolutionary trajectories could inform the development of more precise, niche-targeted therapeutic strategies.

## Intro

Antimicrobial resistance (AMR) is an urgent global threat, with over 2.8 million AMR infections in the United States each year, causing at least 1.27 million deaths worldwide in 2019 (1,2). At the current increasing rate of AMR infections, antimicrobial infections could cause more than 30 million deaths by the year 2050 (3). Although the prevalence of AMR infections is on the rise, the development and approval of novel antibiotic therapies has decreased over the last decade, leading to overuse and the perpetuation of AMR (4). One of the most serious threats for antimicrobial resistance is *P. aeruginosa*, with more than 30,000 hospitalizations and 2,700 deaths per year in the United States due to multidrug-resistant (MDR) infections with nearly $5 billion in associated annual healthcare costs (1,5). *P. aeruginosa* is a versatile opportunistic pathogen that quickly evolves resistance to an array of antimicrobial classes via modulation of membrane permeability and expression of efflux systems, among other adaptations (5–9). Emerging research suggests that metabolic reprogramming may play a pivotal role in the development of AMR (6,7).The link between metabolism and development of resistance presents a significant opportunity for novel therapeutic targeting. However, metabolic modulation in *P. aeruginosa* in the context of the active development of AMR is poorly understood.

Bacteria can evolve resistance over the course of an infection within a host, highlighting metabolic flexibility dependent on environment (8–10). Furthermore, *in vitro* experiments show that bacteria differentially evolve microbial resistance based on the media they are grown in, demonstrating the impact of site-specific metabolites within the host (11). For example, L-alanine is a preferred carbon source for *P. aeruginosa* in the lungs of patients with cystic fibrosis (12). It has been demonstrated that exogenous glucose and alanine restore antibiotic susceptibility in several AMR bacterial strains, including *P. aeruginosa* (13). In *P. aeruginosa*, fumarate and glyoxylate perturb central carbon metabolism in different ways, resulting in altered antibiotic susceptibility phenotypes (7). Furthermore, the nutrient profile in the host environment has been shown to be an important evolutionary driver in AMR development (8).To understand the evolution of and to develop mitigation strategies for AMR, it is critical to clarify the relationship between bacterial metabolism and antimicrobial tolerance and resistance. Investigating how host-specific environments potentiate the development of AMR is a foundational first step to understanding the interaction between bacterial metabolism and the development of resistance.

In addition to diverse metabolic milieu across infection sites, oxygen content can also fluctuate throughout infection. For example, in cystic fibrosis (CF), early lung infections typically occur on the surface of the airways and alveoli, and therefore the microbes exist in a more aerobic environment. Conversely, chronic CF lung infections are enveloped in thick mucus, often further enveloped in a biofilm, significantly reducing oxygen availability (14). These shifts in oxygen availability can drive metabolic switching from aerobic to anaerobic metabolism in bacteria (15). Furthermore, the efficacy of certain antimicrobials is oxygen dependent, causing treatment challenges for infections with fluctuating oxygen availability (16). Consequently, there is a significant need for an antimicrobial that is robust to environmental changes across host infection sites. Determining the mechanistic relationships between aerobic and anaerobic conditions and the development of AMR will allow us to identify metabolic patterns and potential therapeutic strategies.

*P. aeruginosa* is an ideal model organism for studying metabolism and AMR due to its genomic flexibility and metabolic versatility (17). This opportunistic pathogen can utilize a wide range of carbon sources and adapt to diverse environments, allowing it to thrive in nutrient-limited conditions, such as those found in hospital settings and chronic infections. For these reasons it is able to colonize a variety of sites, traditionally the lung, urinary tract, and chronic wounds despite different microenvironments (18). The genetic plasticity of *P. aeruginosa* enables it to rapidly develop resistance to multiple antibiotics regardless of mechanism of action or infection site. *P. aeruginosa* is frequently implicated in MDR infections with some MDR *P. aeruginosa* isolates resistant to nearly all antibiotics, including carbapenems, which are typically considered “last line” antibiotics. Additionally, its well-characterized genome and availability of genetic tools make it a powerful model organism for studying the relationship between metabolism and antibiotic resistance.

In this study, we investigated how distinct host-mimicking environments constrain and shape the evolutionary trajectories of antibiotic resistance in *Pseudomonas aeruginosa*. While traditional studies predominantly evaluate resistance evolution under standardized, nutrient-rich laboratory conditions, we performed long-term experimental evolution in synthetic media specifically formulated to recapitulate the nutrient, chemical, and physical constraints of chronic wound and urinary tract environments, alongside standard Lysogeny Broth (LB) controls. Furthermore, we systematically isolated the specific environmental variables of pH and oxygen tension to decouple global nutritional effects from physiological stress responses during resistance development. Across these diverse conditions, populations were selected for resistance against three clinically vital antibiotics spanning distinct functional classes: piperacillin (PIP), tobramycin (TOB), and ciprofloxacin (CIP). Whole-genome sequencing of strains evolved under contrasting aerobic and anaerobic conditions enabled us to map the genomic and mutational landscapes driving adaptation across these distinct microenvironments.

We show that both the evolutionary rate and the underlying mechanistic pathways of resistance are strictly contingent upon environmental constraints. Under aerobic, nutrient-replete conditions, populations predominantly acquired resistance through canonical target-site and efflux mutations. In contrast, nutrient-limited and anaerobic microenvironments favored distinct suites of genomic mutations, driving modifications in multi-drug efflux systems and metabolic pathways involved in biofilm formation. Importantly, when benchmarked against a large clinical dataset of P. aeruginosa genomic isolates, the mutations identified in our host-mimicking *in vitro* evolutions closely mirrored variant profiles observed in patient infections—confirming that tailored synthetic media capture authentic translational evolutionary trajectories. By establishing a direct link between microenvironmental context, metabolic rewiring, and clinical resistance selection, this work provides a clearer understanding of how *in vivo* conditions dictate evolutionary outcomes, offering critical insights for refining treatment strategies against localized chronic infections.

## Results

### Media composition influences resistance development via nutritional availability

To investigate how the environment influences the development of antibiotic resistance, we evolved *P. aeruginosa* on defined media that represent the host metabolic environments of common infection sites: defined synthetic chronic wound exudate medium (SCWE) (19,20), synthetic human urine media (SHU) (21), and LB media. Using these media, we evolved *P. aeruginosa* resistance to three antibiotics: piperacillin (PIP), tobramycin (TOB) and ciprofloxacin (CIP) (Figure 1A).

**Figure 1.**
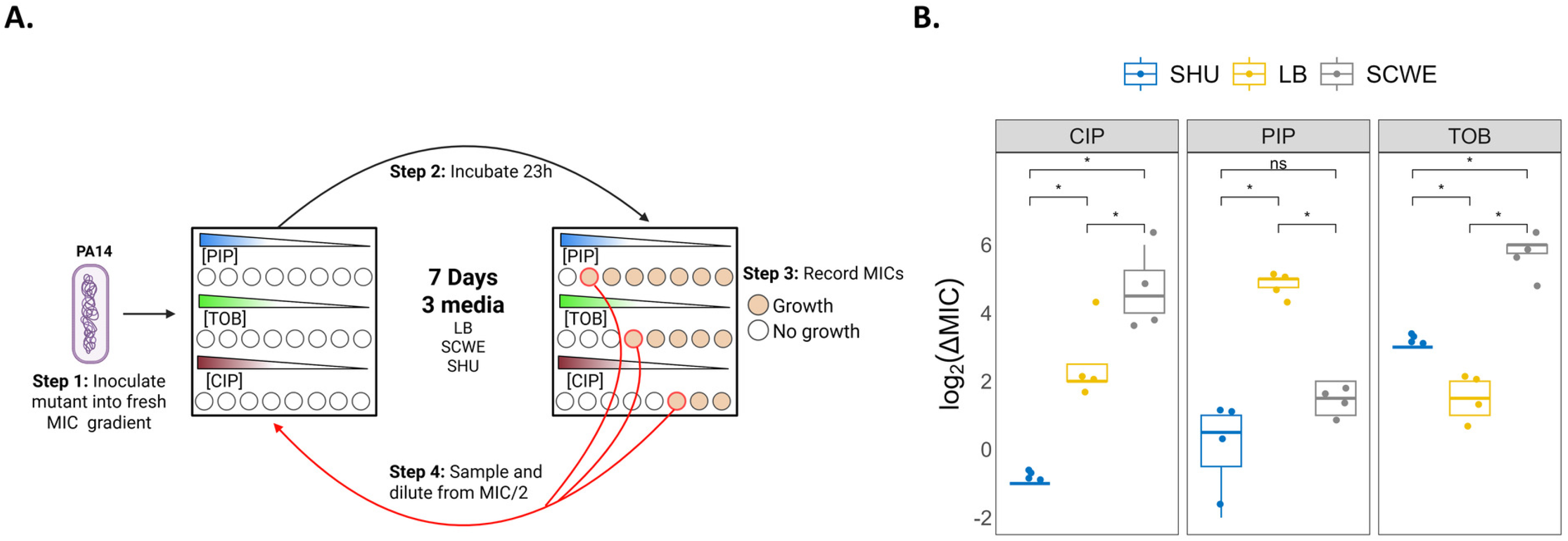
Evolving *P. aeruginosa to* different physiological media. **(A)** Experimental layout for antibiotic-evolution experiment to three different antibiotics: piperacillin, tobramycin and ciprofloxacin, in three different media: LB, Synthetic Chronic Wound Exudate (SCWE) and Synthetic Human Urine (SHU). (B) Changes in the MIC of the PA14 ancestor strain when evolved to each drug in each of the 3 media.

We quantified the log-fold change in MIC between day 7 and day 1 for each condition and compared them within each drug group using a Kruskal-Wallis test with a post hoc Dunn’s test (Figure 1B). We found that for each drug, the media had a significant effect on the rate of evolution. When treated with ciprofloxacin, the cultures exhibited a 5-fold MIC increase in the synthetic chronic wound media, while those in the SHU media experienced a decrease in MIC over time. Interestingly, the media’s effects on the rate of evolution were different for both PIP and CIP. When evolved to PIP, the cultures experienced a 5-fold MIC increase in the LB media compared to the SCWE and SHU media, which both had nearly no increase in MIC. The TOB-evolved isolates experienced nearly a 6-fold increase in MIC in SCWE with minimal MIC increases in the other two.

The SCWE media is designed to mimic a physiological fluid that is rich in amino acids and carbon sources that support robust central metabolism. Previous work has shown that metabolic activity can influence mutation rates and antibiotic tolerance by altering redox balance and DNA repair pathways (16,17,22). The richer nutrient profile of SCWE may have led to the increased metabolism that supports an increased rate of development of resistance to TOB and CIP relative to the other media. In contrast, SHU medium imposes nutrient limitations, particularly in carbon sources, which can restrict bacterial growth and reduce the metabolic activity required for resistance mechanisms to emerge (7,23). This environment may instead select for phenotypes geared toward persistence or survival at low metabolic cost rather than classical high-level resistance (23,24). The lower CIP MIC observed in SHU-evolved populations could reflect fitness trade-offs in which maintaining resistance mutations were too costly under nutrient-restricted conditions.

The differences between LB and the host-mimicking media highlight the importance of studying the development of antibiotic resistance in physiologically relevant settings. LB is a rich, artificial medium that supports rapid growth and may favor resistance mechanisms not traditionally observed *in vivo* (25). Isolates evolved to PIP achieved the highest increase in MIC in LB media compared to the others. Although both LB and SCWE are both nutrient-rich, they produced distinct resistance profiles, indicating that overall carbon-substrate richness alone does not predict the trajectory of resistance evolution. Instead, the specific nutrient composition of the medium must play a critical role.

### Medium pH modulates ciprofloxacin and tobramycin resistance evolution

The physiological media described above are complex and formulated to mimic targeted body sites. The synthetic media used for these evolutions were calibrated to physiologically relevant pH levels: 5.5 for the synthetic human urine media, 7.4 for the SCWE and 7.0 for the LB media. To specifically evaluate how pH affects the development of antibiotic resistance, we also evolved PA14 for 10 days to PIP, CIP and TOB in LB media at a pH of 5.5, 6.5 and 7.5 (**Figure 2**). We found that altering the pH of LB had no significant effect on the development of PIP resistance. In contrast, low pH reduced the rate of resistance adaptation to both CIP and TOB. Ciprofloxacin activity is known to be enhanced under acidic conditions, in part because protonation improves uptake and DNA gyrase targeting, thereby increasing selective pressure and limiting the fixation of resistance mutations (26). Similarly, aminoglycosides such as tobramycin rely on proton motive force for uptake, which can be disrupted under low pH, reducing drug entry into cells and altering the balance between tolerance and resistance (27). The absence of a pH effect on piperacillin resistance suggests that PIP resistance mechanisms, which often involve mutations in *ampC* or penicillin-binding proteins, are less dependent on extracellular pH, and thus evolve similarly across conditions. Although these trends are observed in LB, they do not fully explain the trends seen in the physiological media. For example, the development of TOB resistance was higher in the SHU media (pH 5.5) than in LB media (pH 6.5), suggesting that factors beyond pH are influencing resistance trajectories.

**Figure 2.**
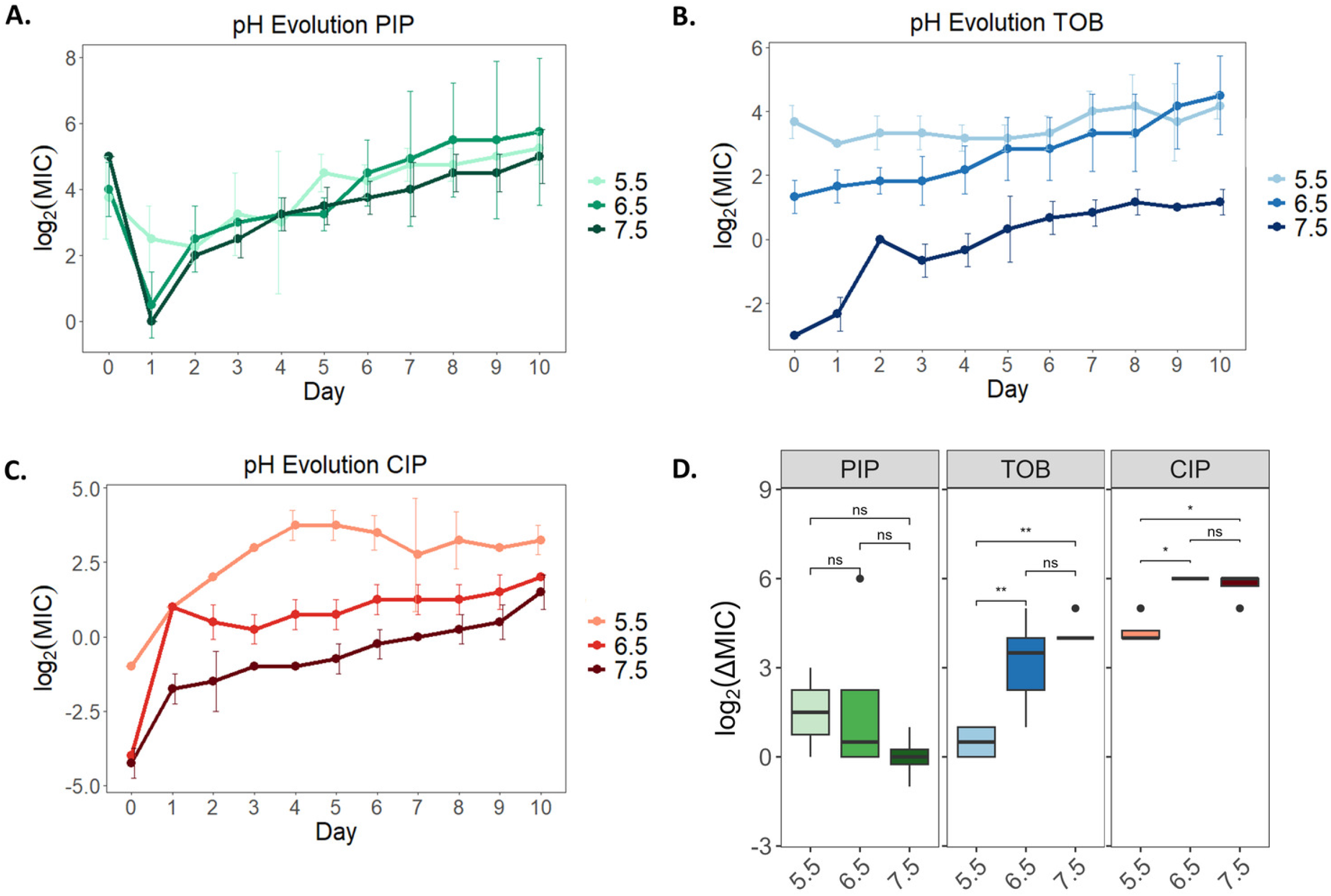
Evolving *P. aeruginosa* to different physiological pH levels. Change in MIC of each of PA14 evolved in LB media to either (A)piperacillin, (B)tobramycin, or (C)ciprofloxacin at different pH levels and their overall MIC fold-change from day 10 compared to day 0 (D).

### Oxygen Availability Shapes Drug-Specific Resistance Evolution Trajectories

To test the role of oxygen on antibiotic resistance development, we evolved PA14 to PIP, TOB and CIP in synthetic cystic fibrosis media (SCFM1)(28) in the presence or absence of oxygen for 15 days (Figure 3). In CIP, the anaerobically evolved strains experienced nearly a 7-fold increase in the MIC while the aerobically evolved strains only had a 2-fold increase in MIC. For PIP and TOB, the aerobically evolved isolates achieved greater increase in their MICs compared to the aerobically evolved isolates. This enhanced resistance in aerobic isolates aligns with previous findings that TOB may have a reduced efficacy in clinical anaerobic conditions (29,30).The reduced efficacy could be due to the reduced entry of antimicrobials into anaerobically grown cells because transfer of certain drugs like TOB is dependent on membrane potential. The use of nitrate as a terminal electron acceptor instead of oxygen may lower the membrane potential, resulting in reduced entry to the cell. Furthermore, reduced entry may not pose the same evolutionary pressure as the levels of TOB in aerobic conditions. CIP enters the cell via diffusion, but resulting antibiotic lethality may be related to the generation of reactive oxygen species (31), which would be decreased in anaerobic conditions. This decrease in reactive oxygen species could protect the cell from oxidative stress-mediated cell death. PIP has also been shown to stimulate increased production of reactive oxygen species, and this production was reduced in the presence of biofilm (32). The presence of biofilm may have limited cell access to oxygen. In our previous mutant evolutions, there were large genomic deletions in the region of the genome containing *hmgA* (33). The deletion of *hmgA* leads to the pyomelanin hyperproduction, which can contribute to AMR (34).The increased oxidative stress induced by PIP could be beneficial for conferring resistance by destabilizing the genome and allowing for mutagenesis via the SOS-response pathways (35). Our sequencing of the PIP-evolved isolates also contained these large genomic deletions, but they only appeared in the aerobically evolved isolates.

**Figure 3.**
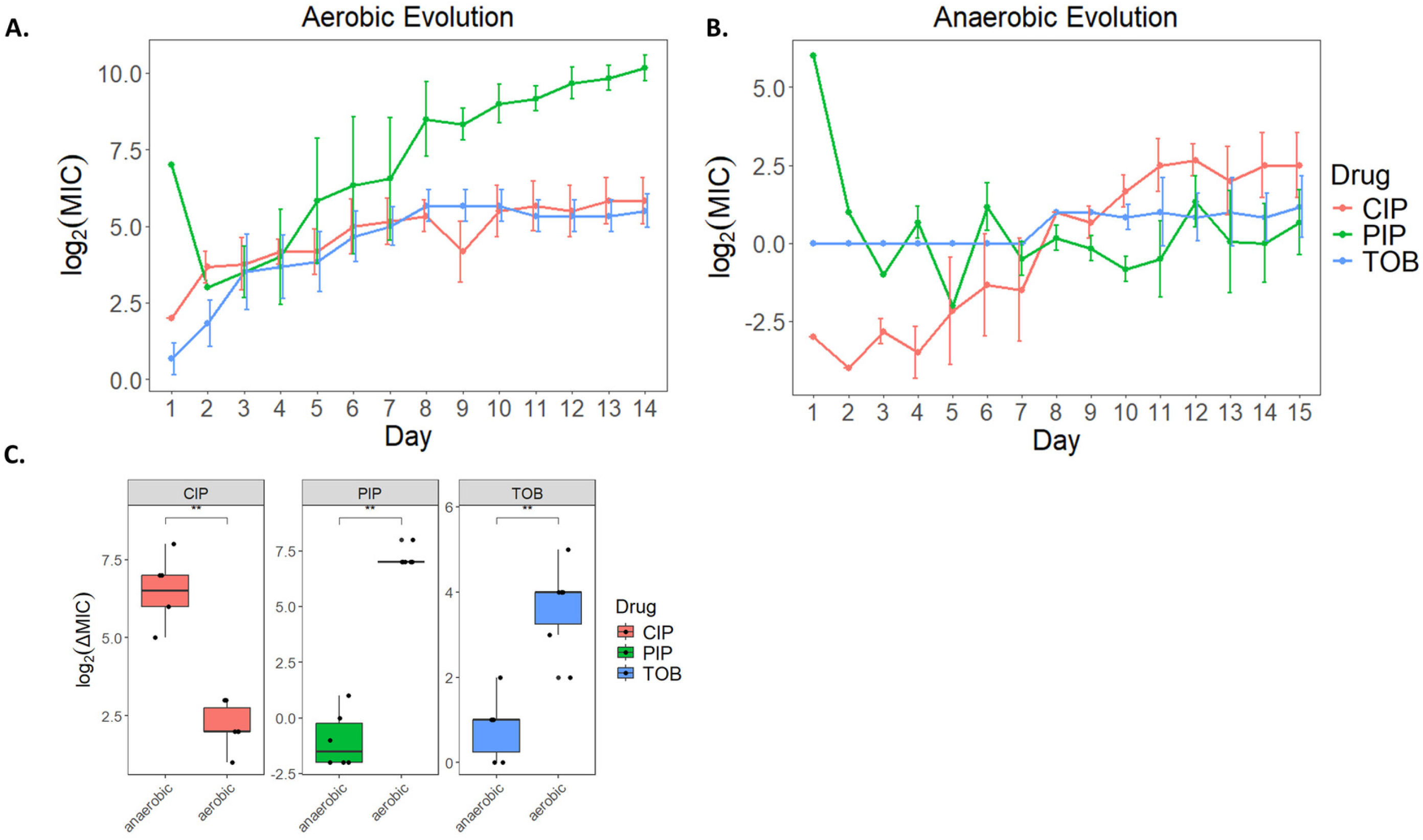
Evolving P. aeruginosa in the presence and absence of oxygen. Change in MIC of each of PA14 evolved in Synthetic Cystic Fibrosis Media (SCFMl) supplemented with nitrates to piperacillin, tobramycin or ciprofloxacin in **(A)** aerobic or **(B)** anaerobic conditions. **(C)** The log2-fold-change in MIC between the last day and the first day of evolution in each condition for each drug.

To test how evolved isolates behave when transferred to a different physiological environment after evolution in another environment, we transferred the strains evolved aerobically to anaerobic conditions to assess MICs without oxygen (Figure 4). We also tested the MICs of the anaerobically-evolved strains in aerobic conditions. When transitioned from aerobic-to-anaerobic growth, the MICs of most PIP-evolved strains decreased. This trend was reversed when anaerobically-evolved strains to PIP were switched to aerobic conditions. The loss of piperacillin resistance under anaerobic conditions may reflect the dependence of β-lactam resistance mechanisms on oxygen-linked metabolic activity. For instance, mutations in *ampC* regulation or penicillin-binding proteins may not provide the same advantage when anaerobic metabolism reduces cell wall biosynthesis rates or alters redox balance (36). Additionally, the phenazine redox recycling operon is contained in the regions affected by the large genomic deletions in our aerobic PIP-evolved isolates. Phenazines help maintain the intracellular redox homeostasis in anaerobic conditions by serving as electron sinks (37), and deleting this operon may create a severe redox bottleneck, halting fermentation capabilities and slowing anaerobic growth.

**Figure 4.**
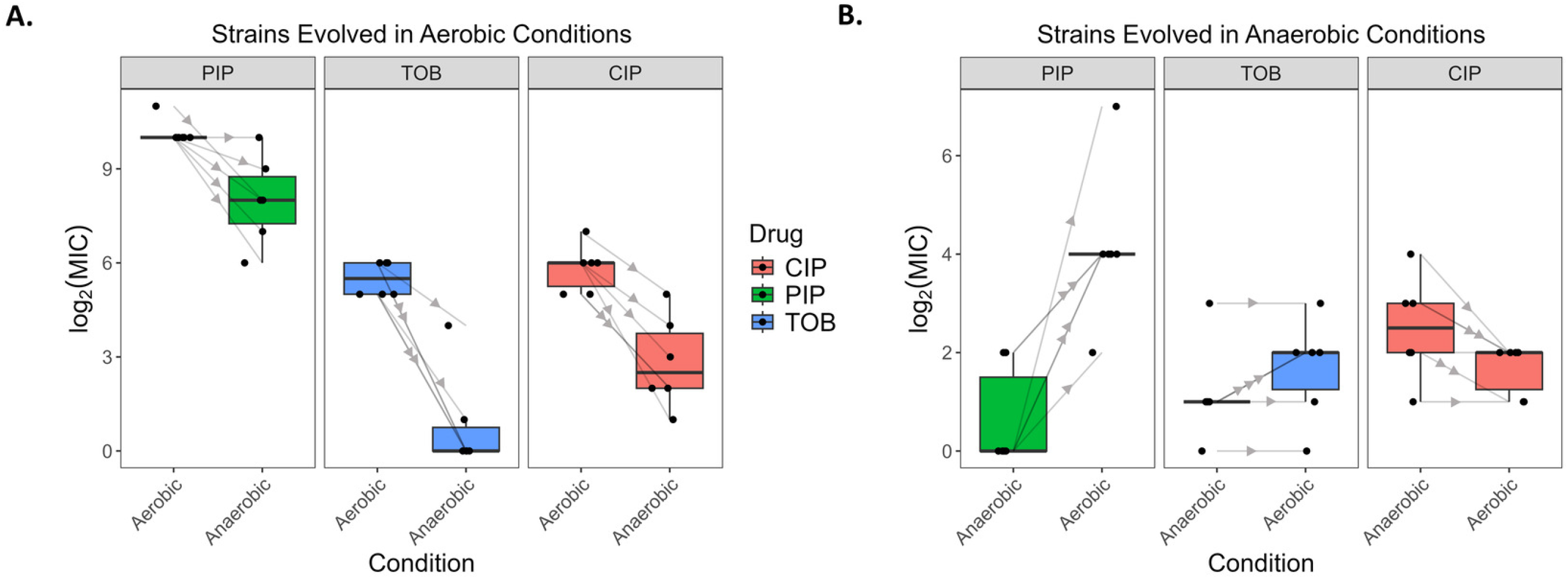
Assessing the change in MIC after evolution to aerobic and anaerobic conditions. MICs to piperacillin, tobramycin and ciprofloxacin after the final day of evolution to (A) aerobic conditions and (B) anaerobic conditions. Each arrow corresponds to a biological replicate.

Tobramycin-evolved isolates experienced a much sharper decrease in MIC when switched from aerobic to anaerobic conditions. Aminoglycoside resistance is closely tied to uptake, which requires proton motive force (PMF) and is usually decreased in anaerobic conditions. Therefore, isolates in anaerobic conditions should have reduced tobramycin entry into the cell. One possible explanation is that certain resistance mechanisms such as ribosomal modification may impose a metabolic cost that, when coupled with the stress of switching to anaerobic conditions, limits the ability of the bacteria maintain homeostasis. Furthermore, the presence mutations specific to aerobically-evolved isolates (**Figure 5**) further suggests that there are trade-offs between resistance acquisition and metabolic flexibility, consistent with the high energetic cost of aminoglycoside resistance pathways in *P. aeruginosa* (38).

**Figure 5.**
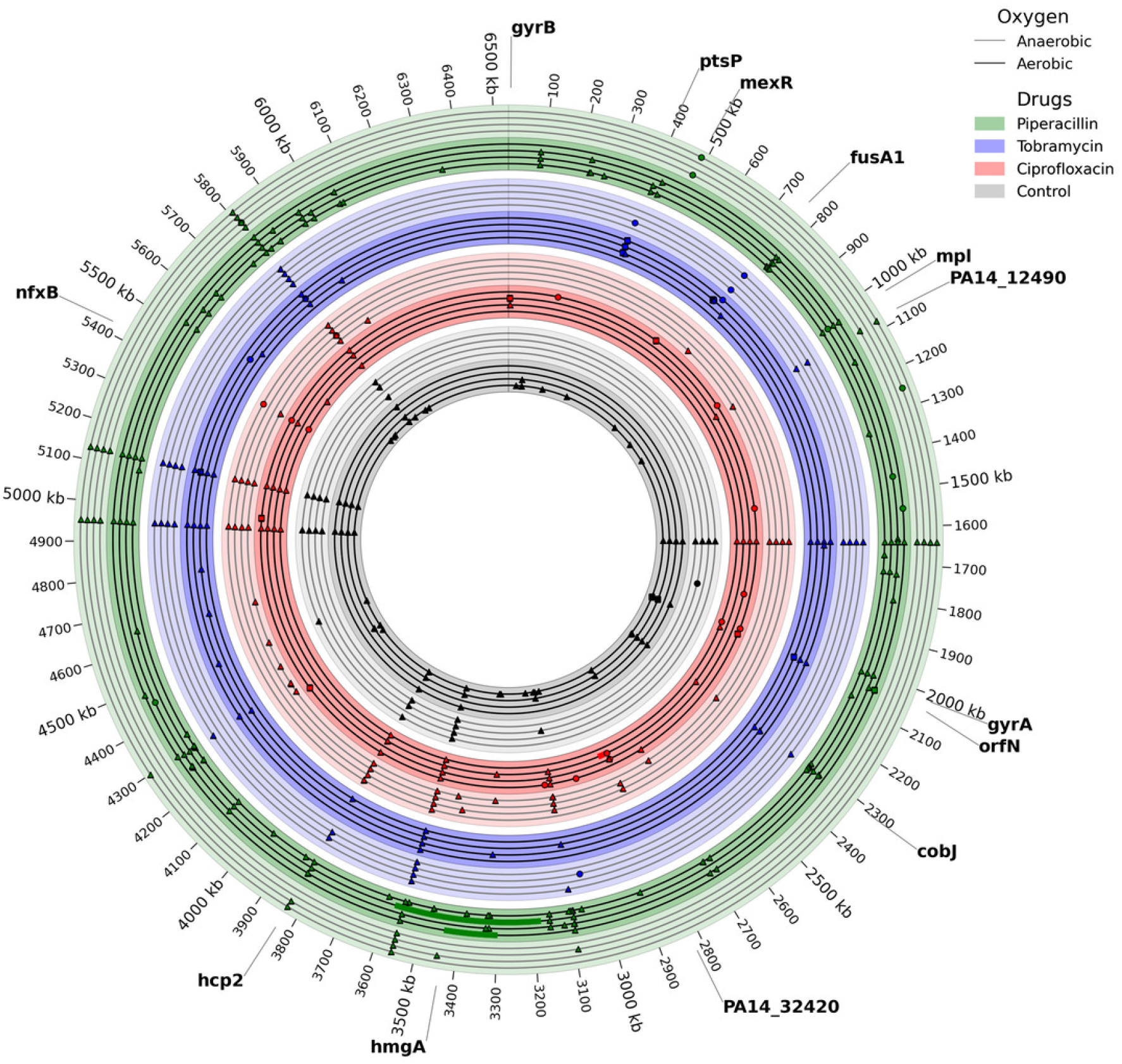
Mutations in the anaerobically and aerobically evolved isolates. Circles represent SNPs, triangles represent small deletions(<lkbp), squares represent insertions, and bars represent large deletions (>Ikbp). Lighter shade genome circles correspond to replicates evolved anaerobically, and darker shades correspond to replicates evolved aerobically.

The CIP-evolved isolates exhibited modest decreases in MIC when shifted between aerobic and anaerobic conditions. Isolates that evolved resistance aerobically showed a slight decrease in MIC when transferred to anaerobic growth, while isolates that evolved anaerobically displayed a similar slight decrease when shifted to aerobic growth. These results suggest that CIP resistance mechanisms selected under one oxygen condition remain largely effective under the other, though with a minor reduction in efficacy. This stability is consistent with the common resistance strategies for fluoroquinolones in *P. aeruginosa*, such as target site mutations in *gyrA* and *parC* or overexpression of efflux systems like MexEF-OprN, which function across oxygen environments (39).

### Whole genome sequencing reveals differing resistance mutations between aerobic and anaerobic conditions

To identify the genetic changes underlying these resistance phenotypes, we sequenced the genomes of isolates evolved under both aerobic and anaerobic conditions, as well as the ancestral strain prior to evolution. We observed differences in both the number of mutations and types of mutations when comparing the strains evolved aerobically to those evolved anaerobically for each drug. The strains evolved aerobically to PIP had an average of 37 mutations while those evolved anaerobically only had an average of just over 8 mutations (Table S1/figure 5). This difference may be because the strains grown aerobically could reach a higher density than those grown anaerobically. Although the strains grown both aerobically and anaerobically had the same number of generations evolved, they had a different number of cells dividing in each generation, which can lead to a difference in mutation rate. Additionally, aerobic metabolism may lead to increased reactive oxygen species (ROS) relative to the strains evolved anaerobically, leading to DNA damage and elevated mutation rates (22,40).

The strains evolved aerobically to PIP contained common evolutionary strategies of resistance. Two of the aerobic PIP-evolved strains contained large genomic deletions of 141kb and 401kb in length. We observed similar deletions in a previous evolution study (33). These deletions both span *hmgA*, a gene that encodes for homogentisate 1,2-dioxygenase which, when disrupted, leads to pyomelanin hyperproduction, which can contribute to AMR (34). All of the aerobically-evolved PIP strains had a mutation in *mpl,* which encodes for an important enzyme in peptidoglycan recycling. *mpl* disruption has been shown to activate β-lactamase production (41) thus facilitating increased resistance to β-lactams. None of the anaerobically-evolved strains contained a mutation in *mpl*. Inactivation of *mpl* derepresses *ampC* beta-lactamase expression by interrupting peptidoglycan recycling (42). Under oxygen-deprived conditions, the metabolic cost of constitutively overexpressing *ampC* may be energetically unsustainable given the reduced ATP yield of anaerobic respiration. Moreover, altered peptidoglycan turnover and cell envelope stress under hypoxia may reduce the selective advantage of *mpl* loss, driving anaerobic populations to favor alternative resistance mechanisms—such as multidrug efflux pump regulation and biofilm-associated metabolic remodeling—that provide piperacillin tolerance without compromising metabolic fitness.

The strains evolved anaerobically to PIP also contained some unique resistance mutations apart from those observed in the aerobic strains. Two of the anaerobic strains contained mutations in *hcp2*, along with one of the aerobic strains. *hcp2* is involved in the type VI secretion system, which is traditionally associated with bacterial competition and has been shown to play a role in biofilm formation (43), which may aid in the development of resistance. Additionally, only strains evolved anaerobically to PIP contained mutations in PA14_12490 and *mexR.* Mutation in *mexR* results in an increased resistance profile because it results in the reduced repression of the *mexAB-oprM* multidrug efflux operon (44). PA14_12490 is a putative protein that likely encodes for AMP Nucleosidase, which is involved in purine metabolism. Purine metabolism has been shown to influence ci-di-GMP and resulting biofilm formation (45), which may play a role in protecting from drug penetration. Collectively, these results demonstrate that anaerobic adaptation comes from both classical resistance mechanisms and by impacting biofilm formation. These adaptations appear to be unique to anaerobic evolution, highlighting the importance of the environmental niche in the development of resistance.

The CIP-evolved isolates also contained differential mutations depending on the presence of oxygen. The aerobic isolates evolved an average of 14 mutations where the anaerobically-evolved isolates evolved an average of just over 12 mutations. Additionally, the aerobic isolates all had either a mutation in *gyrA* or *gyrB*, which encode for subunits of DNA gyrase. These mutations are a primary mechanism of resistance to fluoroquinolones. None of the anaerobically evolved isolates had either of these mutations. The CIP-evolved anaerobic and aerobic strains also shared some common mutations such as PA14_32420, which encodes for *mexS,* a putative oxidoreductase. Mutational inactivation of *mexS* is a primary driver of *mexEF-oprN* overexpression, typically occurring through the functional activation of the transcriptional regulator MexT (46,47). Mutations in *mexS* were found in two isolates from both the anaerobically and aerobically evolved strains to CIP. Similarly, both the aerobic and anaerobic strains contained mutations in *nfxB*, leading to the up-regulation of *mexCD-oprJ* expression and, consequently, increased resistance to fluoroquinolone antibiotics (48). The shared occurrence of mutations in *mexS* and *nfxB* highlights the importance of multidrug efflux regulation as a mechanism of adaptation that is independent of oxygen availability. However, the absence of *gyrA/gyrB* mutations in anaerobically evolved isolates indicates that selective pressures differ substantially between aerobic and anaerobic environments, potentially shaping distinct evolutionary trajectories toward ciprofloxacin resistance.

TOB-evolved isolates experienced similar evolution patterns to that of CIP. The aerobically-evolved isolates had an average of over 31 mutations compared to just 8 mutations in the anaerobically-evolved isolates. Anaerobic isolates evolved to TOB more often had mutations in the *fusA1* gene. The *fusA1* gene encodes elongation factor G (EF-G1A), which plays a critical role in protein synthesis during translation elongation and ribosome recycling. It has been associated with aminoglycoside resistance and is often observed in clinical isolates from chronic infections of patients with CF (49). Aerobic strains frequently contained mutations in *cobJ* and *orfN,* which encode for a methyltransferase involved in B12 synthesis and O-antigen biosynthesis. Although not commonly associated with antibiotic resistance, these mutations can affect central metabolism and surface antigen structure, possibly contributing to modulated antibiotic susceptibility (50). Aerobic strains also frequently contained a mutation in the *ptsP* gene, which encodes for the phosphoenolpyruvate phosphotransferase protein. This protein is part of the nitrogen phosphotransferase system and has been identified to be a target of TOB resistance for *P. aeruginosa,* although its exact mechanism of resistance is unknown (51).

Two anaerobic strains also had mutations in PA14_12490, similar to the anaerobic PIP-evolved isolates. Interestingly, only anaerobically-evolved strains contained this mutation. Two of the strains that were passaged anaerobically without the presence of any drug also contained this mutation. Since this gene is likely involved in biofilm formation, it may be an adaptation to thrive in anaerobic conditions rather than an adaptive response to the drug. It will require further testing to determine this gene’s role in resistance.

### Validation of Experimental Resistance Mutations in a Clinical P. aeruginosa Genome Cohort

To evaluate whether resistance-associated mutations identified in our lab-evolved strains translate to clinical isolates, we analyzed a database of 414 fully sequenced *P. aeruginosa* clinical genomes with data on resistance to TOB and CIP. Isolates were systematically benchmarked against the PA14 reference strain to evaluate MIC distributions for CIP and TOB as a function of variation across 7 commonly mutated candidate target genes (*gyrA, gyrB, cobJ, ptsP, fusA,* PA14_32420, and *nfxB*). To resolve variant-specific phenotypic impacts, empirical cumulative distribution functions (ECDFs) were constructed for the top statistically significant locus per gene, comparing wild-type (WT) and mutant populations across both antimicrobial classes (Figure 6). Although these are clinical isolates, the isolates that did not differ from the PA14 reference genome at the specific locus being observed will be referred to as “wild-type.”

**Figure 6.**
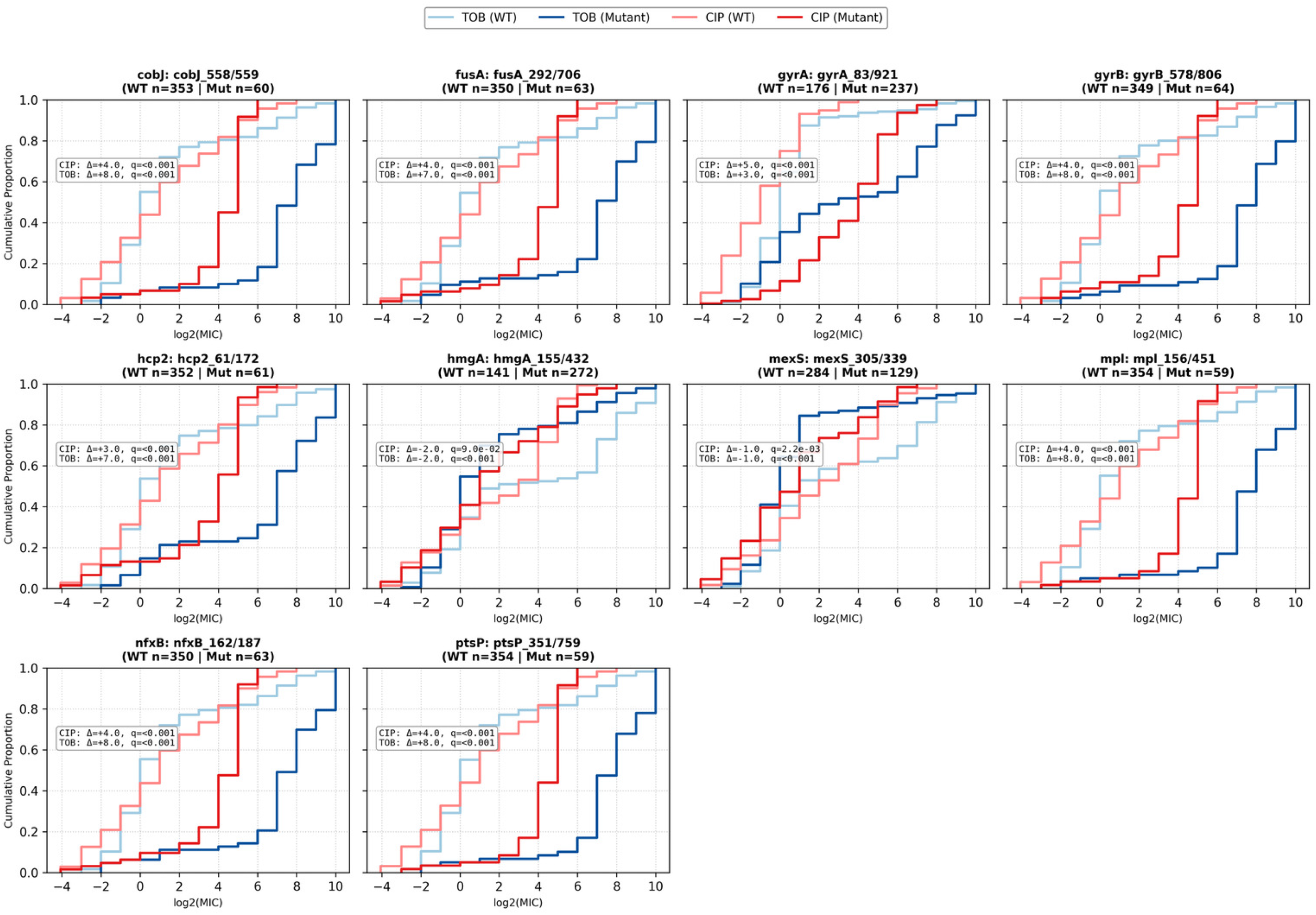
Empirical cumulative distribution functions of MICs for top significant resistance loci across Tobramycin and Ciprofloxacin. Subplots compare the MIC distributions log_2_-transformed between Wild Type (light shades) and Mutant (dark shades) isolate populations for the most statistically significant variant per candidate gene (q < 0.05). Sample sizes for WT and mutant groups are indicated in panel titles. Inset text boxes display the median log. and false discovery rate -adjusted q-value (Mann-Whitney U test) for each drug independently. A rightward shift of mutant curves relative to wild type indicates increased antibiotic resistance.

This clinical translation revealed striking disparities in phenotypic impact: while substitutions at the 83rd amino acid position of *gyrA* (e.g., T83I) drove a 5 log_2_-fold increase in CIP MIC (32-fold linear shift) alongside a co-associated 3 log_2_-fold increase in TOB MIC, variants in PA14_32420 produced no meaningful phenotypic shift, inducing a slight decrease in the MIC of isolates to CIP and TOB. The remaining candidate loci exhibited nearly identical ECDF curve trajectories, consistently producing an approximate 4 log_2_-fold increase in CIP MIC and an 8 log_2_-fold increase in TOB MIC relative to wild-type isolates.

The uniformity among the remaining ECDF profiles highlights a fundamental divergence between experimental evolution and clinical epidemiology. In controlled laboratory selections, mutations arise sequentially in response to isolated antibiotic pressures, allowing for the precise isolation of single-gene fitness effects. In contrast, clinical isolates are shaped by complex host environments, prolonged multi-drug regimens, and clonal transmission. The near-identical phenotypic shifts observed across distinct gene targets strongly indicate that these mutations do not act as independent, isolated resistance mechanisms in the clinic. Instead, they represent genomic co-occurrence, a phenomenon where high-level multi-drug resistant isolates harbor multiple target mutations simultaneously (52).

This clustering is likely amplified by clonal expansion and regional sampling biases, where circulating hospital lineages accumulate pan-resistant genotype profiles over time (53,54). Consequently, while classical target mutations like *gyrA* Thr83 function as primary drivers of resistance, many secondary loci appearing statistically significant in univariate clinical screenings are passenger mutations in highly resistant genetic backgrounds. Disentangling true causal drivers from genetic linkage artifacts remains a crucial step when translating laboratory-evolved resistance mechanisms into clinical diagnostic models. Additionally, the fact that anaerobic-specific (*fusA*) and dual-condition (*nfxB*) mutations co-occur alongside aerobic drivers (*gyrA*) in clinical isolates suggests that circulating hospital lineages sequentially adapt to shifting anatomical niches over time. Additionally, it is likely that these isolates are from chronic infections, which have been treated with multiple drug classes, and have had sufficient time to build up genetic resistance to several drugs in different environments.

## Discussion

This study aims to demonstrate that the metabolic environment strongly shapes the evolutionary trajectories of *P. aeruginosa* under antibiotic pressure. By comparing evolution across host-mimicking synthetic media, pH variation, and oxygen availability, we reveal how distinct physiological conditions influence both the rate and mechanism of resistance development. Our findings highlight that the presence of oxygen in the environment plays a pivotal role in determining whether resistance emerges through canonical mutational pathways or via alternative adaptations such as efflux regulation and biofilm-associated traits.

One of the key findings of this work is that the composition of the growth medium substantially altered resistance outcomes. SCWE promoted rapid adaptation to both tobramycin and ciprofloxacin, whereas SHU constrained resistance evolution. These results support the notion that nutrient-rich environments with diverse carbon sources enhance potential bacterial metabolism, thereby increasing opportunities for mutagenesis and selection of resistance. In contrast, nutrient-limited conditions such as SHU may impose strong fitness constraints, selecting instead for tolerance or persistence strategies rather than actively developing higher levels of resistance. Such dynamics could help explain why chronic wound infections are often associated with multidrug-resistant *P. aeruginosa*, whereas urinary tract infections more commonly present with persistent but not necessarily highly resistant populations (55).

Unlike high-level resistance, which relies on inherited genetic mutations that elevate the MIC, tolerance and persistence are non-heritable physiological states characterized by reduced metabolic activity that allow bacteria to survive lethal antibiotic exposure without altering MIC. These distinct evolutionary trajectories suggest clear therapeutic implications. In urinary tract infections, where nutrient limitation favors persistent or tolerant populations rather than high-level genetic resistance, increasing antibiotic dosage is often ineffective against dormant cells; instead, therapeutic approaches should emphasize combination therapies or metabolic adjuvants that “awaken” persister cells to restore antibiotic susceptibility. Conversely, in chronic wound environments where abundant nutrients accelerate mutagenesis and selection for high-level resistance, simply ramping up drug concentration risks driving further genetic adaptation. These infections may be better targeted by addressing the specific metabolic and physical features of the wound microenvironment, such as disrupting biofilms or altering nutrient availability to constrain evolutionary pathways alongside antimicrobial therapy.

Our findings also emphasize that pH influences resistance in a drug-specific manner. Low pH slowed the adaptation of CIP and TOB resistance, consistent with enhanced ciprofloxacin activity under acidic conditions and reduced aminoglycoside uptake due to impaired proton motive force. In contrast, PIP resistance evolved independently of pH, underscoring that β-lactam resistance mechanisms such as *ampC* derepression or alterations in penicillin-binding proteins are less sensitive to extracellular proton concentration. Importantly, however, the pH effects observed in LB medium did not fully explain the differences between SHU and SCWE, highlighting that the nutrient environment exerts an influence beyond pH alone.

Oxygen availability emerged as another major axis governing resistance trajectories. CIP resistance evolved more readily under anaerobic conditions, whereas PIP and TOB resistance increased rapidly under aerobic growth. Aminoglycoside uptake relies on an active electron transport chain to generate a proton motive force (56). Under anaerobiosis, reduced membrane potential impairs drug entry, decreasing selection pressure for target mutations. For fluoroquinolones, lethality is partly mediated by reactive oxygen species (ROS) downstream of target binding (31). Under anaerobic conditions, reduced ROS-mediated cell death permits sub-lethal mutant survival, accelerating resistance emergence. Furthermore, the rapid loss of piperacillin resistance when aerobically evolved strains were transitioned to anaerobiosis suggests that *β*-lactam resistance mechanisms depend on oxygen-linked metabolic processes, such as peptidoglycan turnover or redox-dependent cell wall regulation (57).

Whole-genome sequencing further illuminated the distinct mutational strategies employed under different environments. Aerobic evolution was associated with higher mutational burdens, likely reflecting both higher cell densities and increased ROS-driven mutagenesis. Large genomic deletions encompassing *hmgA* were common in aerobically evolved piperacillin isolates, consistent with previous studies linking pyomelanin production to resistance and genomic instability (58). Conversely, anaerobic evolution frequently involved mutations in genes linked to efflux regulation (*mexR, mexS, nfxB*) and biofilm-associated functions (*hcp2*, PA14_12490). These findings suggest that while aerobic conditions select for canonical target-site and cell-wall associated resistance mechanisms, anaerobic conditions favor adaptive strategies that remodel biofilm physiology and efflux capacity, potentially offering broader survival benefits in low-oxygen infection niches such as cystic fibrosis sputum.

Finally, drug-specific patterns of mutation were evident. Ciprofloxacin resistance under aerobic conditions consistently involved *gyrA/gyrB* mutations, the classical mechanism of fluoroquinolone resistance (59), whereas anaerobic adaptation favored efflux-related mutations in *mexS* and *nfxB*. Tobramycin resistance in anaerobic populations often arose through *fusA1* mutations, mirroring mutations observed in cystic fibrosis clinical isolates (49), whereas aerobic isolates more frequently carried mutations in *cobJ* and *orfN*, genes not classically associated with resistance but plausibly affecting metabolism and surface structure. The recurrence of PA14_12490 mutations specifically under anaerobic evolution, even in the absence of antibiotic, suggests that some genetic adaptations may represent environment-specific metabolic optimization rather than direct drug responses.

This work shows that the environment not only modulates the rate of resistance development but also likely constrains the underlying genetic mechanisms. This context-dependence has significant clinical implications. Laboratory studies conducted in rich artificial media such as LB may overestimate certain resistance trajectories while overlooking physiologically relevant pathways that arise in host-like environments. Moreover, our findings suggest that strategies to mitigate resistance could be informed by the infection niche. For instance, interventions targeting efflux activity or biofilm physiology may be particularly effective in anaerobic or nutrient-limited environments.

Translating our laboratory-evolved data to a database of 414 clinical *P. aeruginosa* genomes revealed striking evolutionary convergence. Variants selected in the lab under aerobic (*gyrA, gyrB, cobJ, ptsP*), anaerobic (*fusA*), and dual conditions (*nfxB, PA14_32420*) were all also present in clinical isolates. This result indicates that over the course of human infections, *P. aeruginosa* adapts across both aerobic and anaerobic microenvironments.

However, translating these findings to the clinic highlights a fundamental difference in evolutionary dynamics. While laboratory experiments isolate single-gene effects under fixed oxygen levels, clinical populations navigate complex spatial oxygen gradients and long-term, multi-drug exposure (60). The uniform MIC shifts observed across diverse clinical targets demonstrate that these mutations do not act as independent resistance mechanisms in patients. Instead, they reflect genomic co-occurrence: multi-drug resistant clinical clades sequentially accumulate mutations across both aerobic and anaerobic infection stages over time.

Consequently, while lab-evolved anaerobic adaptations like *fusA1* represent authentic responses to low-oxygen niches (frequently observed in cystic fibrosis isolates; (49)), their strong statistical association with high-level resistance in clinical surveys is largely an artifact of genetic linkage within these heavily mutated, pan-resistant lineages. This result explains why these target loci share near-identical ECDF profiles across the clinical cohort: rather than reflecting independent resistance mechanisms, they trace the same subset of multi-drug resistant strains that sequentially acquired tolerance and resistance to both CIP and TOB across shifting aerobic and anaerobic infection niches.

The evolutionary pathways to antibiotic resistance in *P. aeruginosa* are tightly constrained by the surrounding physical and metabolic environment. By recapitulating host-relevant conditions, including oxygen availability, nutrient limitations, and pH variation, we demonstrated that distinct microenvironments select for specialized, niche-specific resistance mechanisms. Crucially, the target variants identified in our laboratory evolution experiments directly mirror mutations observed in clinical populations, validating that local physiological pressures *in vivo* drive predictable evolutionary trajectories. Recognizing that clinical resistance often reflects an accumulation of niche-adapted variants within multi-drug-resistant lineages allows us to better anticipate how *P. aeruginosa* adapts during chronic infection. Ultimately, decoding the complex intersection of bacterial metabolism, environmental constraints, and resistance evolution will deepen our understanding of pathogen adaptation and guide more precise, niche-targeted therapeutic strategies.

## Methods

### Experimental study design

For both the media and pH evolution experiments, four independent replicates were evolved. For the aerobic-anaerobic evolution, six independent replicates were evolved. The number of replicates was chosen to balance both the statistical power and technical feasibility of serial propagations and daily media preparation. For the media and pH evolution, a period of 7 and 10 days was selected because the similarities and differences of the drug and media-specific effects were readily apparent at that point. For the aerobic-anaerobic evolution, a period of 15 days was selected because the resistance levels of the evolved lineages began to stabilize at that point.

### Media, growth conditions, and antibiotics

MIC plates were made daily using the broth microdilution method with the standard 2-fold dilution series (61). LB was used as the growth medium for all experiments unless otherwise specified (1% tryptone, 0.5% yeast extract, 1% NaCl). Antibiotics tested include piperacillin sodium, tobramycin, and ciprofloxacin HCl (Sigma). Aliquots of 1 mg/ml and 10 mg/ml antibiotic stocks were made by diluting the antibiotic powders in LB and were stored at −20°C. New frozen drug aliquots were used on a daily basis.

### Adaptive laboratory evolution in physiological media and pH

This evolution protocol is based on a previously published protocol (33). A frozen stock of *P. aeruginosa* PA14 was streaked on an LB agar plate, and a single colony was inoculated into 5 ml of each media type (LB, SCWE, SHU), which was then grown overnight at 37°C, shaking at 150 RPM. The physiological media formulations come from previous publications (synthetic chronic wound exudate (19,20), synthetic human urine media SHU (21), and the preparation guidelines are outlined in Table S2. This antibiotic-susceptible culture, denoted as the Day 0 Ancestor, was diluted to an OD600 of 0.01 and then inoculated into three identical MIC plates consisting of concentration gradients of piperacillin, tobramycin, and ciprofloxacin in their respective media. A sample of the ancestor was saved in 25% glycerol and stored at −80°C. The three MIC plates were used to serially propagate cultures evolved to LB media, piperacillin, and tobramycin, with four biological replicates per condition. Wells for growth control (media + culture) and sterility control (media) were included in each MIC plate. MIC plates were placed in a plastic container (to prevent evaporation) and incubated at 37°C with shaking at 125 RPM (Thermo Scientific MaxQ 4000). MIC plates were incubated daily for approximately 23 hours.

Growth was defined as an OD600 > 0.1 after background subtraction and was measured at 23 hours using the Tecan Infinite M200 Pro. We defined the minimum inhibitory concentration (MIC) as the lowest antibiotic concentration tested that did not show growth in each medium and drug combination. The cultures from the highest concentration that showed growth (MIC/2) were inoculated into a fresh new gradient of the drug. This was repeated each day for seven days. For adaptation to the media alone, cultures were passaged from the growth control well that contained only the media without any drug. For each culture to be passaged, the culture was first diluted by a factor of 1/225 in fresh LB (e.g., 8 μl of the culture was diluted in 1.8 ml of media), which was then inoculated in fresh piperacillin, tobramycin, or ciprofloxacin drug gradients in the MIC plate for the new day. Wells of the MIC plate thus contained 100 μl of double the final concentration of the antibiotic and 100 μl of the diluted culture. Hence, the cultures were diluted by a total factor of 1/450 daily. Daily samples were saved in 25% glycerol and stored at −80°C.

The same protocol was used to evolve PA14 to PIP, CIP and TOB at different pH levels as described above. Shortly after autoclaving the LB media, the pH was adjusted to either 5.5, 6.5 or 7.5 and filter sterilized using a 0.2µm Nalgene Rapid Flow bottle top filter.

### Adaptive laboratory evolution in aerobic and anaerobic conditions

For the aerobic-anaerobic evolution protocol, PA14 was streaked from frozen stock onto a LB agar plate and incubated overnight at 37°C. A single colony was inoculated into 5ml of Synthbiome Synthetic Cystic Fibrosis Media (SCFM1) (28) and grown overnight in a shaking incubator at 37°C and 150RPM. We chose SCFM1 because it contains all the metabolic components of cystic fibrosis sputum, without the DNA, mucin, N-Acetylglucosamine and dioleoylphosphatidylcholine. This reformulation allowed for our evolutions to focus on the metabolic adaptations that occur during evolution. Also, this medium is transparent, which is ideal for our spectrophotometric-based growth assessments. 10 ml of the culture was then put into two 5ml tubes of SCFM1 with 10mM of potassium nitrate to support anaerobic growth. One tube was placed in a regular incubator and the other was placed in a Bactron300 anaerobic chamber, which allows for less than 4.7 parts per million oxygen. Both cultures were allowed to grow overnight at 37°C shaking at 150RPM. These cultures were then diluted to an OD600 of 0.01, and placed into MIC plates for the evolution procedure as described above. All the passaging for the aerobic cultures was performed in a standard biosafety cabinet and all the passaging for the anaerobic cultures was performed in the anaerobic chamber. All OD_600_ readings for this experiment were performed with a Cerillo Alto (Cerillo, Inc.) so that the reads could be performed in the anaerobic chamber. Each day, new aliquots of media were placed in the anaerobic chamber to reduce for 23 hours for use the next day.

### Whole genome sequencing

Frozen samples were streaked on LB agar plates and incubated overnight at 37°C in the evolved oxygen condition of the isolate. A single colony was inoculated into LB+10mM of nitrate and placed in a shaking incubator overnight at 37°C and 150 RPM in the evolved oxygen condition of the isolate. The culture was then centrifuged to a pellet and the liquid was aspirated. The pellet was then frozen at -80°C and sent to Seqcenter (Pittsburgh, PA) for whole genome sequencing.

The processing done at Seqcenter are as follows. Illumina sequencing libraries were prepared using the tagmentation-based and PCR-based Illumina DNA Prep kit and custom IDT 10bp unique dual indices (UDI) with a target insert size of 280 bp. No additional DNA fragmentation or size selection steps were performed. Illumina sequencing was performed on an Illumina NovaSeq X Plus sequencer in one or more multiplexed shared-flow-cell runs, producing 2x151bp paired-end reads. Demultiplexing, quality control and adapter trimming was performed with bcl-convert (v4.2.4).

### Variant Identification in Laboratory-Evolved Isolates

To identify *de novo* mutations arising during experimental evolution, raw paired-end Illumina sequencing reads from lab-evolved *P. aeruginosa* isolates were first quality-filtered and trimmed using fastp to remove low-quality bases and adapter sequences (62). Mutations were called using breseq (63) using the standard consensus mode settings. All reported mutations were inspected by viewing the read alignments in IGV and the breseq output files, and only mutations with 80% frequency or greater were counted. Mutations were called against the reference *P*. *aeruginosa* PA14 genome (NC_008463.1, (64)). We also sequenced our own PA14 ancestor strain and used the breseq-subtract command to ensure that we only captured mutations that occurred during the lab-evolution.

### Clinical Genome Collection and Variant Calling

To translate experimental findings to clinical populations, a cohort of 414 fully sequenced *P. aeruginosa* clinical genomes with paired MIC metadata for CIP and TOB was retrieved from a previously published study (65). Raw paired-end fastq reads were quality-filtered and adapter-trimmed using Trimmomatic to ensure high-quality sequence input (Q > 20)(66). Trimmed reads were input directly into Snippy (v4.6.0)(67) using the *P. aeruginosa* UCBPP-PA14 reference genome (NC_008463.1, (64)). Snippy managed read mapping via BWA-MEM (68) and variant calling via FreeBayes (69).

### Statistics

Reported MIC values were log_2_-transformed. ECDFs were generated comparing log_2_(MIC) distributions between wild-type and mutant populations for the top locus per candidate gene across both CIP and TOB. An isolate referred to as “wild-type” is an isolate that does not have a mutational difference from the PA14 reference genome in the specific gene or portion of the gene being investigated. Phenotypic distribution differences between wild-type and mutant groups were evaluated using two-sided Mann-Whitney U tests. p-values were adjusted for multiple testing using the Benjamini-Hochberg procedure, with statistical significance established at an FDR q-value < 0.05.

## Data and code availability

The raw sequencing datasets supporting the conclusions of this article are available in the NCBI Sequence Read Archive (SRA) repository under accession PRJNA1531176 (https://www.ncbi.nlm.nih.gov/sra/PRJNA1531176). Code to process the data and generate all figures can be found on at (https://github.com/jmf7ak/media_evolution_paper)

## Additional affiliations

Jason Papin is a co-founder of Cerillo, Inc.(A Hamilton company) which manufactures the Alto plate reader used to generate some of the reported data.

